# Enhanced 2D structured illumination microscopy: super-resolution with optical sectioning and reduced reconstruction artifacts

**DOI:** 10.64898/2026.02.26.708245

**Authors:** Sylvia Marie Steinecker, Henning Ortkrass, Jasmin Celine Schürstedt-Seher, Annika Kiel, Angela Kralemann-Köhler, Jan Schulte am Esch, Thomas Huser, Marcel Müller

**Affiliations:** Biomolecular Photonics Research Group, Faculty of Physics, Bielefeld University, Bielefeld, Germany; Department of General and Visceral Surgery – Liver and Tumor Biology, Medical Faculty OWL, Bielefeld University, Bielefeld, Germany; Department of General and Visceral Surgery, University Hospital OWL, Campus Bielefeld-Bethel, Germany

## Abstract

Structured Illumination Microscopy (SIM) provides imaging with spatial super-resolution, as well as optical sectioning capability, without relying on specialized fluorescent dyes. 2D and 3D variants of this method exist, but most bespoke implementations are 2D-SIM, because it is easier to realize and modify than 3D-SIM. 2D-SIM systems, however, often experience reconstruction artifacts from out-of-focus contributions, especially when pushing for high lateral spatial resolution in thicker samples. We present *enhanced 2D-SIM*, an approach to 2D-SIM where both, coarse patterns optimized for removing out-of-focus background, and fine patterns optimized for resolution improvement beyond the diffraction limit are used. In combination, this achieves 2D-SIM reconstructions with high contrast, spatial super-resolution, and significantly reduced reconstruction artifacts. We present the theoretical framework of this technique, and provide enhanced 2D-SIM imaging results of liver sinusoidal endothelial cells stained with fluorophores emitting at visible and near-infrared wavelengths. Quantitative comparisons of power spectral distribution and image resolution are demonstrated.

## 1 Introduction

Structured illumination fluorescence microscopy (SIM) is currently being utilized in two main applications: Optical sectioning SIM (OS-SIM) uses coarse illumination patterns and allows to computationally separate in-focus signals from out-of-focus background. It allows for the high imaging speeds inherent to widefield-based microscopy methods, while aiming for background reduction similar to laser scanning confocal imaging methods [1, 2, 3]. Super-resolution SIM (SR-SIM), on the other hand, uses fine illumination patterns, typically with a pattern spacing close to the diffraction limit, to modulate the fluorescent signal with a known structure of high spatial frequency. In an algorithmic image processing step, this modulation can be used to down-shift information into the pass band of the microscope, and thus yield previously inaccessible high-frequency information about the sample. As a result, SR-SIM improves the spatial resolution compared to Abbe-limited widefield imaging [4, 3].

SR-SIM can be implemented with different pattern structures, the most popular being two-beam and three-beam SIM, where “beams” refers to the number of laser illumination beams that interfere in the sample plane. In two-beam SIM, only a lateral modulation of the pattern is achieved, while in 3-beam SIM, a central beam path is added and additional axial modulation of the pattern is observed. Based on their typical applications, 2-beam SIM [5, 6] is often referred to as 2D-SIM, while 3-beam SIM is referred to as 3D-SIM [7].

While 3D-SIM is a very capable technique and the most common commercially available implementation of SIM, the method also has a clear drawback, as experienced by many groups that set out to build custom SIM systems: Implementing 3-beam interference with high pattern contrast and frequency is a far greater engineering challenge than building a robust 2-beam SIM system. Bespoke 2D SR-SIM imaging systems are used [8, 9, 10], for example, when cost is a factor [11, 12], or where new avenues for applying SIM imaging are explored [13]. Thus, in practice, 2-beam systems remain a popular choice, with many robust systems having been designed, published, and being readily available [14].

Here, we present an enhancement to 2D-SIM that can be implemented on many custom 2D-SIM systems. It overcomes one of the main drawbacks of classical 2D-SIM systems, the trade-off that is currently needed between aiming either for maximum resolution enhancement or the suppression of out-of-focus background. We first set out to provide the theoretical framework of how the different SIM modalities work and where enhanced 2D-SIM fits in, and then present experimental data acquired with different 2D-SIM modalities.

### 1.1 The missing cone problem of the 3D optical transfer function

A key point in understanding how SIM interacts with out-of-focus structures is examining its 3-dimensional optical transfer function (OTF). It is well known that the 3D OTFs of widefield microscopes [15, 16, 17, 18] feature the so-called *missing cone* [17, 19], i.e. the fact that at low lateral spatial frequencies, there is no axial support in the OTF of a widefield microscope (see Fig. 1(a)). Moving to SIM, in the simplified ideal case, the SIM reconstruction process yields a copy of the OTF at any spatial frequency present in the SIM illumination pattern - scaled with the modulation depth of that modulation, the so-called SIM bands. The angle at which the beams interfere, which sets how fine the SIM pattern can become, is the same as the numerical aperture that defines the diffraction-limited resolution of the system, the Abbe limit. This yields the often-quoted factor of 2× resolution improvement in SIM imaging, though there are some caveats: In practice, to achieve a pattern with high phase stability and modulation depth despite the imperfections which are present in any real-world optical system, super-resolution SIM is typically not performed at the finest pattern physically possible, but at 80% to 90% of the minimum pattern spacing. Also, while the pattern spacing is set by the excitation wavelength, the detection OTF shape and extent is set by the Stokes-shifted emission wavelength.

**Figure 1:**
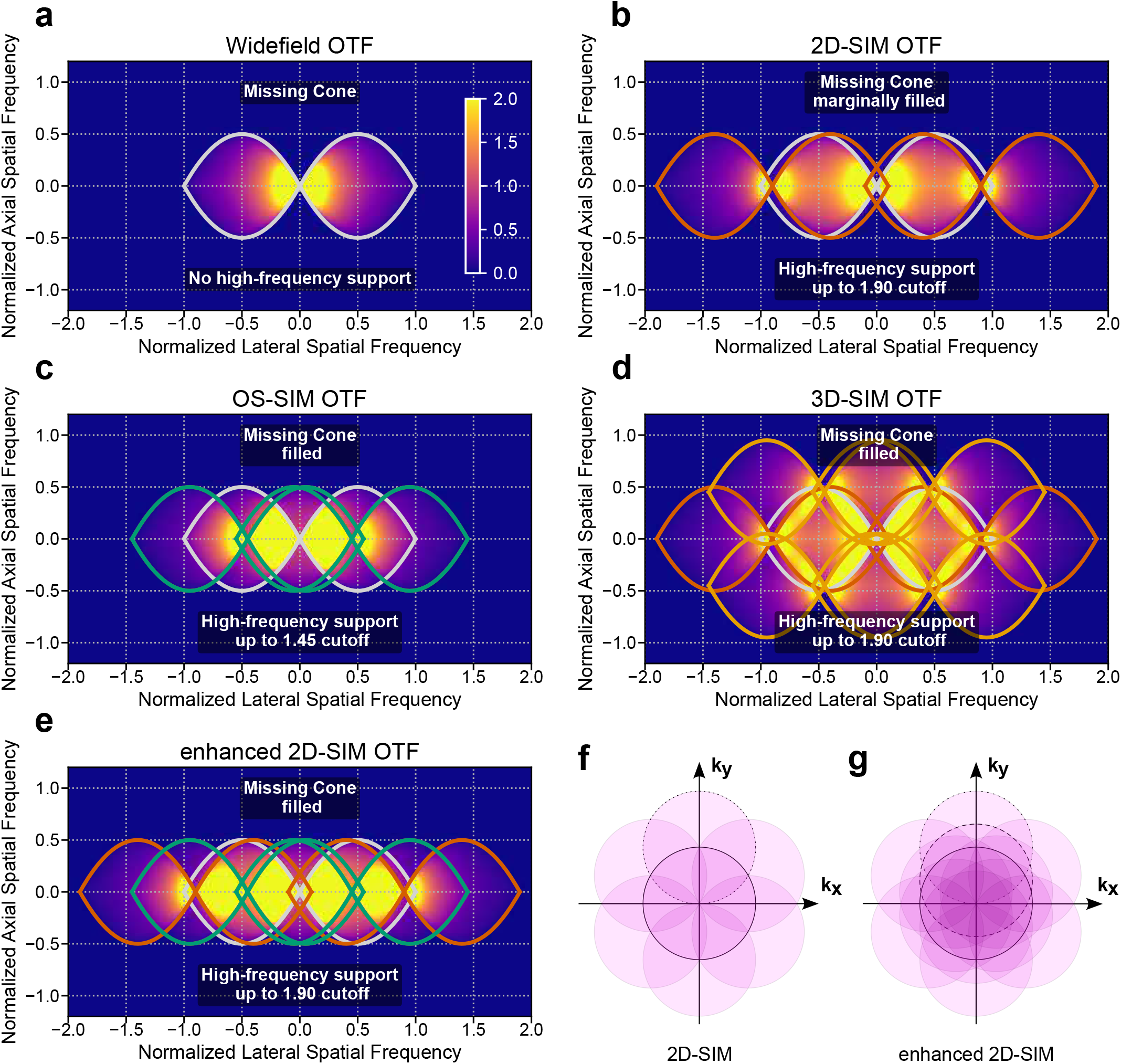
Comparison of optical transfer functions for widefield and structured illumination microscopy variants. 3D optical transfer functions (OTFs) are displayed in (a–e) for widefield (a), 2D-SIM (b), optical sectioning SIM (c), 3D-SIM (d) and enhanced 2D-SIM microscopy (e). The color scale in (a–e) shows the combined normalized OTF intensity, while outlines show the widefield (gray), coarse pattern (green), base frequency (3-beam SIM, yellow) and fine pattern 2-beam (orange) or first harmonic 3-beam (orange) support. The coarse pattern for OS-SIM, 2D-SE-SIM as well as the base frequency for 3D-SIM is set at 45% of the diffraction limit, while the fine pattern for 2D-SIM and 2D-SE-SIM is set at 90% of the diffraction limit, which coincides with the first harmonic of 3D-SIM. Modulation contrast is set at 1 for all 2D-SIM variants, and at 50% central beam intensity, corresponding to a modulation of 1.13 for the low and 0.8 for the high frequency band for 3D-SIM. In simplified terms, due to the SIM reconstruction process, copies of the OTF emerge at any spatial frequency in the pattern, scaled with the modulation depth of that pattern frequency. These also shift the so-called *missing cone*, the fact that at low lateral spatial frequencies, the OTF shows no axial support. In classical 2D-SIM, where the OTF copies are shifted far out by a very fine pattern, the missing cones can barely be filled by information from other SIM bands, which is the main reason why 2D-SIM is prone to out-of-focus artifacts. In all other SIM modalities (OS-SIM, 3D-SIM and also the enhanced 2D-SIM proposed here) the missing cone can be filled with information from higher SIM bands. Plots (a)–(e) show lateral–axial cross sections along the SIM pattern orientation, while for (near) isotropic coverage, additional rotations of the pattern around their central symmetry axis (f,g) are required.

For 2D-SIM with a fine illumination structure, as required for high lateral resolution enhancement, a problem becomes obvious: By pushing the OTF far out along the lateral axis, the missing cones of the SIM band and the widefield overlap and are not filled (see Fig. 1(b)). This explains the problem leading to out-of-focus background artifacts which are typical in 2D-SIM reconstructions, and heavily limit for which samples 2D-SIM can be used.

There are two established solutions to this problem: When the main goal of SIM imaging is optical sectioning and not lateral resolution enhancement, a much coarser SIM pattern can be used, requiring only 3 lateral pattern shifts at a single angle [1, 2]. This reduces how far the SIM bands are shifted out, and yields an overlap of the transfer functions in the missing cone region (see Fig. 1(c)). OS-SIM therefore yields robust suppression of out-of-focus contributions. When OS-SIM data are acquired with nine raw images (3 angles with 3 phase shifts per angle) and are subjected to SR-SIM image reconstruction, a minimal improvement in lateral resolution can be obtained [20]. Alternatively, the illumination structure employed by 3-beam SIM [7] has a very convenient property: Due to the presence of the third, central beam the lateral modulation is composed of a base frequency at or slightly below half of the frequency cutoff, and a first harmonic close to or at the cutoff of the optical system. In the reconstruction step, this allows to fully fill the frequency space, and thus also the missing cone (see Fig. 1(d)). By using 15 raw images, 3D-SIM extends both lateral and axial frequency support (close to) 2× the resolution limit, and avoids out-of-focus artifacts.

A third option is explored here: By using a 2-beam SIM system where the pattern spacing can rapidly be switched between a coarse and a fine pattern, the optical sectioning capability of OS-SIM can be combined with the high lateral resolution improvement achieved by classical 2D-SIM. Instead of acquiring both SIM bands simultaneously as in 3D-SIM, the coarse and fine pattern are applied sequentially and then combined in the reconstruction step. The resulting effective transfer function is the combination of OS-SIM and 2D-SIM, it fills the missing cone and extends lateral resolution (see Fig. 1(e)). Using 3 pattern orientations (see Fig. 1(f,g)) to reconstruct near-isotropically resolved images, this method requires a minimum of 18 instead of 15 raw images. However, this is somewhat mitigated by the modulation depth achievable for both modalities: In 2D-SIM (ideal case), both patterns modulate at 100%. In 3D-SIM, however, modulation can be shifted between base frequency and the first harmonic by varying the central beam intensity. In the OTFs presented here, we assume the central beam to be at 50%, which yields a modulation of 1.13× for the first and 0.8× for the second SIM band.

### 1.2 Single-slice reconstruction and effective OTFs

The classical reconstruction process for 3D-SIM [7] images works on 3D stacks of image data, and implicitly fills the 3D missing cone of the optical transfer function as part of the reconstruction process. However, and crucially, the capability of suppressing out-of-focus background contributions does not depend on full 3D imaging, but can also be applied to single-slice data, as long as there is overlap of OTFs that fill the regions of the missing cones. Now, with these 2D datasets, a specific processing protocol is needed: The so-called OTF attenuation [21, 22] is used to artificially dampen information at spatial frequencies where the missing cone is present in the 3D OTF. For each band, the attenuation is applied at the spatial frequency defined by the respective SIM illumination pattern from which the band originates. The attenuation is often described as notch-filtering, which is accurate when looking only at a single SIM band. However, in the combined reconstruction, it is a notch-shaped re-weighting of frequency contributions around the missing cones, as information filtered out in one band (due to the missing cone) is filled again by information from an overlapping band that does not feature the missing cone at the same spatial frequency.

As a result, slightly different effective optical transfer functions are obtained for the different modalities, which are summarized in Fig. 2. The OTFs of the different widefield imaging modalities without attenuation are shown in Fig. 2(a). Fig. 2(b) compares the effective transfer functions for a flat sample, where there is no need to fill the missing cone and thus to attenuate the OTF. Fig. 2(c) shows the attenuated OTFs for comparison. Fig. 2(d) uses a typical OTF attenuation around the missing cone of each band. The high pattern frequency is set at 90% of the OTF support, which leaves some overlap to fill the missing cone, but classical 2D-SIM still shows clear dips in its OTF at zero and at pattern frequency, which show up as patterned artifacts in the SIM reconstruction process. All other OTFs (OS-SIM, enhanced 2D-SIM, 3D-SIM) also show dips where the missing cone needs to be compensated, but in practice they are shallow enough to still provide a reliably, artifact-free reconstruction.

**Figure 2:**
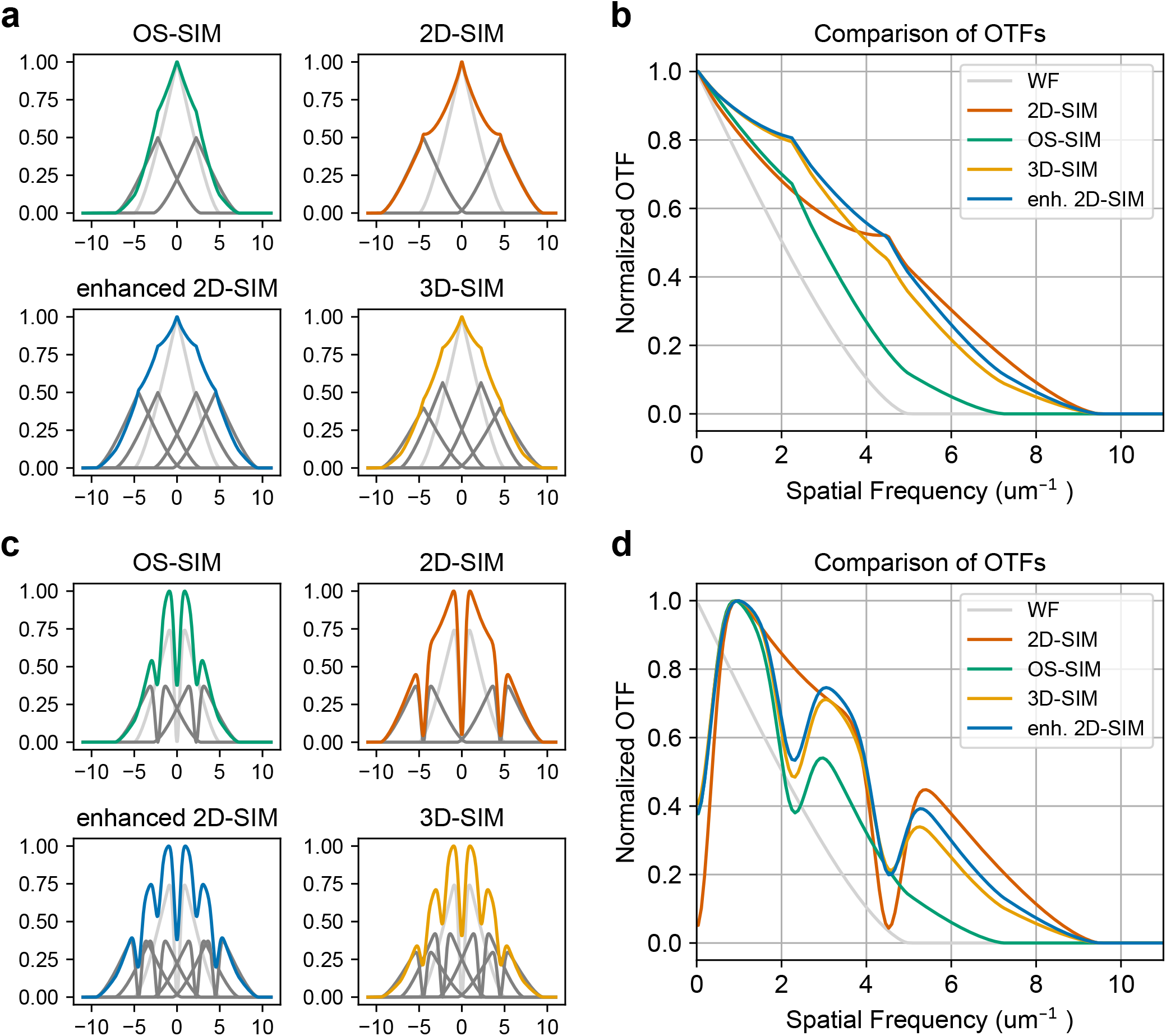
Effective optical transfer functions (OTFs) for different SIM variants. In panels (a) and (c), the widefield contribution is shown in light grey, the SIM band contributions in dark gray, and the resulting effective OTF in color. Panels (b) and (d) show the effective OTF for all modalities for comparison. The top row panels (a) and (b) show the OTF without any attenuation. The bottom row panels (c) and (d) show an attenuated version of the OTF, where the missing cone is filled by re-weighting, and, thus, out-of-focus artifacts are suppressed. Noticeably, there are steep dips in the attenuated 2D OTF at DC and 0.45 µm^−1^, which explains the poor performance of classical 2D-SIM when out-of-focus light is present. The OTFs are computed at 200 nm resolution limit (1.25 NA, 500 nm wavelength), with 444 nm coarse and 222 nm fine pattern spacing, a modulation of 1 for 2D-SIM and a central beam intensity of 50%, corresponding to modulations of 1.13 and 0.8 for 3D-SIM. The attenuation is set at 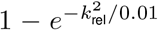 or specifically 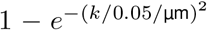, with the distance measured relative to the center of the corresponding SIM band, and k_rel_ given in units of the OTF cutoff.

## 2 Methods

To experimentally demonstrate and verify enhanced 2D-SIM, we have upgraded two bespoke 2-beam SIM systems with the ability to rapidly switch between coarse and fine SIM patterns, and acquired data-sets at different excitation and emission wavelengths. One of these systems operates at standard visible light wavelengths, while the other works in the near-infrared (NIR) spectral range, where there are no readily available solutions for 3D-SIM imaging.

### 2.1 Optical systems and imaging modalities

The SIM systems used in this work have been previously described in Refs. [13, 23]. Compared to these implementations, the main modification introduced here is the replacement of the segmented (“pizza”-shaped) half-wave plate (HWP) with a custom-built HWP rotator that provides greater flexibility in polarization control and allows for coarser pattern spacing. A schematic of the optical setup is shown in Fig. 3(a).

**Figure 3:**
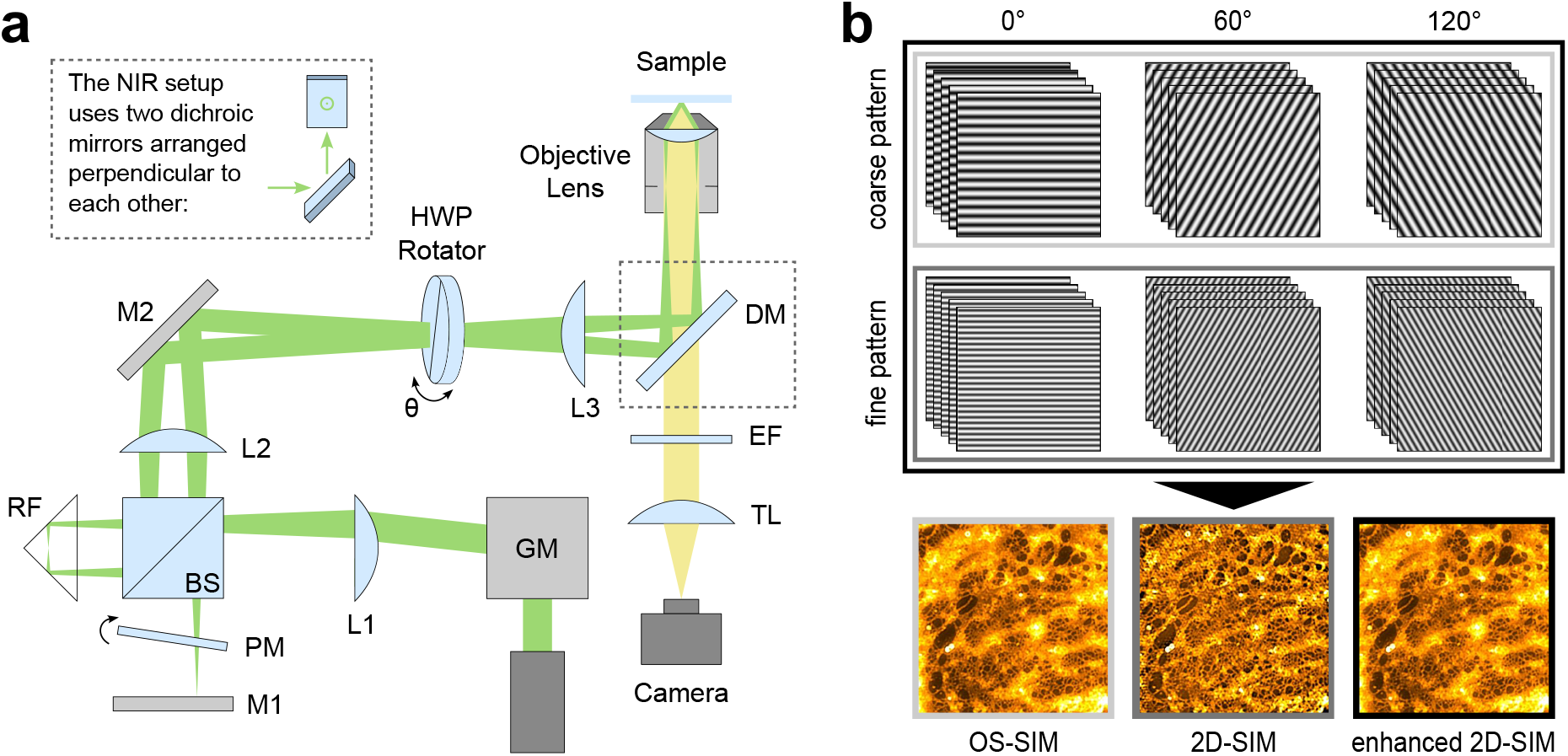
Setup schematics and imaging modalities for OS-SIM, 2D-SIM and enhanced 2D-SIM. (a) The SIM setups are based on a modified Michelson interferometer. Excitation laser light is deflected by a pair of galvanometric mirrors (GM) and focused by lens L1. A non-polarizing 50/50 beamsplitter cube (BS) divides the beam into two interferometer arms: one containing a retroreflector (RF) and the other a phase modulator (PM) and mirror M1 for phase shifting. The two resulting divergent beams, symmetrically displaced with respect to the optical axis, are relayed by a telescope formed by lens L2 and L3 and directed via mirror M2 and a polarization-maintaining dichroic mirror (DM) into the back focal plane of the objective lens. A custom-built half-wave plate rotator ensures s-polarization in the sample plane for all illumination angles. Fluorescence emission is filtered by an emission filter (EF), collected by the tube lens (TL) and imaged onto an sCMOS camera. In the NIR setup, two non-polarization-maintaining dichroic mirrors arranged orthogonally are used to compensate for polarization distortions. (b) Conventional SIM data sets with three illumination angles and five phase steps are acquired using both a coarse and fine illumination pattern, enabling three imaging modalities. Reconstruction using only the coarse pattern yields OS-SIM images, providing efficient background suppression and a moderate resolution improvement compared to widefield imaging. Reconstruction using only the fine pattern corresponds to 2D-SIM, offering higher spatial resolution but often with increased susceptibility to reconstruction artifacts. The enhanced 2D-SIM approach combines raw data from both, coarse and fine patterns during reconstruction, integrating the background rejection capability of OS-SIM with the resolution enhancement of 2D-SIM while reducing artifacts caused by out-of-focus fluorescence.

Briefly, both systems are based on a modified Michelson interferometer [23, 24, 25] in which one arm contains a retroreflector (RF), while the second arm serves as a phase-shifting arm incorporating a phase modulator (PM) and a mirror (M1). The excitation beam is steered by a pair of galvanometer mirrors (GM) positioned in the back focal plane of lens L1 and subsequently focused by L1. This configuration causes the beam to propagate parallel to the optical axis with a lateral offset from the optical axis as it passes through a non-polarizing 50/50 beamsplitter cube (BS). While the RF reflects the beam symmetrically about the optical axis, phase modulation is achieved in the second arm by using a tilting glass plate mounted on a modified galvanometer mirror scanner (where the mirror is replaced by a planoparallel glass plate). Tilting this plate in the beam introduces controlled changes in the optical path length, thereby generating phase shifts. The two beams are relayed by a 4f-telescope formed by lenses L2 and L3 and redirected by mirror M2 and a polarization-maintaining dichroic mirror (DM) before being focused into the back focal plane of the objective lens. For the NIR configuration a polarization-maintaining dichroic mirror was not available at the operating wavelength. Therefore, two perpendicularly arranged dichroic mirrors were used to compensate for polarization distortions [12, 13]. This optical arrangement allows for the free adjustment of the illumination pattern spacing, orientation, and phase solely by controlling the GM and PM positions. Polarization is controlled by using the custom-built HWP rotator, which consists of a half-wave plate mounted in a ball bearing that is rotated by a stepper motor. The rotator is synchronized with the rest of the system to ensure s-polarized illumination in the sample plane for each SIM orientation, maximizing modulation depth [26]. Compared to the previously used segmented HWP, the rotator offers greater flexibility by enabling arbitrary pattern spacings and SIM angles without dead zones from “pizza”-segment edges. The HWP rotator is placed in a conjugate pupil plane between L2 and L3 to utilize its clear aperture most efficiently. Fluorescence signal is collected in epi-detection by the objective lens, transmitted through the DM, filtered by an emission filter (EF) and imaged onto the camera by a tube lens (TL). The VIS system uses an objective lens with NA = 1.5 and an excitation laser with λ = 561 nm, while the NIR system has an objective lens with NA = 1.45 and a laser source with λ = 785 nm.

For this work, 15 SIM raw images (three illumination angles with five phase steps each) were acquired for both a coarse and a fine illumination pattern. In principle, the total number of images can be reduced to 18 by using the minimum of nine SIM raw images per pattern spacing [6, 4]. We chose the increased number of phase steps to further reduce artifacts in the image reconstruction process. Raw data were processed using the Fiji plugin fairSIM [27], which uses the Gustafsson reconstruction approach for super-resolution SIM [6]. Reconstructing only the data acquired with the coarse illumination pattern results in an OS-SIM image, which provides effective suppression of out-of-focus background but only a moderate improvement in lateral resolution compared to widefield imaging. Reconstruction based solely on the fine illumination pattern yields a conventional 2D-SIM image with enhanced spatial resolution, while exhibiting significant reconstruction artifacts, particularly in bright image areas. Our enhanced 2D-SIM reconstruction method combines both data sets by reconstructing the coarse and fine pattern raw images. In fairSIM, this can easily be done by selecting six illumination angles instead of the standard three. By combining both pattern spacings, enhanced 2D-SIM links the background rejection capability of OS-SIM with the resolution enhancement of 2D-SIM. In addition, the inclusion of the coarse pattern introduces increased overlap in the mid-frequency range, improving frequency support and reconstruction stability, which further reduces reconstruction artifacts compared to conventional 2D-SIM (see Section 1). The principle of the different imaging modalities is illustrated in Fig. 3(b). The modified Michelson-type interferometer used in this work is particularly well suited for such novel imaging modalities, as it allows flexible control over illumination angles and pattern spacings, making it effortless to switch between them during imaging. However, as only two pattern spacings are needed, typical implementations of SIM using spatial light modulators or digital micromirror devices could also be adapted for this imaging method.

### 2.2 Sample preparation

To compare the established imaging modalities OS-SIM and 2D-SIM with the enhanced 2D-SIM approach, samples with fixed liver sinusoidal endothelial cells (LSECs) were prepared. Similar to the procedure described previously [13], LSECs were obtained from RjHan:NMRI mice housed by the Animal Facility at Bielefeld University and cell isolation was carried out according to a previously published protocol [28]. The liver was perfused immediately after sacrificing the animal, enzymatically digested and the resulting cell suspension was subjected to a series of centrifugation steps. LSECs were subsequently purified from the non-parenchymal cell fraction using CD146-coated magnetic beads (MACS, Miltenyi). Cells were seeded at a density of 2.5 × 10^6^ cells/ml in EGM medium supplemented with 2% fetal bovine serum onto #1.5 borosilicate glass coverslips coated with 0.2 mg/ml fibronectin. Cultures were maintained at 37°C in a humidified incubator with 5% CO_2_ and 5% O_2_ until use.

For VIS membrane imaging, cells were incubated with BioTracker 555 Orange Cytoplasmic Membrane Dye (SCT107, Merck KGaA) diluted 1:142 in CO_2_-independent medium (Gibco, ThermoFisher) for 20 min at 37 °C. For NIR imaging, BioTracker NIR790 Cytoplasmic Membrane Dye (SCT115, Merck KGaA) was used at a dilution of 1:1000. After staining, VIS samples were washed three times with fresh CO_2_-independent medium and fixed with 4% PFA in PBS for 10 min at 37 °C. NIR-fluorescent samples were washed in the same manner and fixed using 3% glyoxal in PBS for 30 min at room temperature [29]. Both samples were rinsed three times with PBS prior to imaging. VIS samples were mounted using Fluoroshield (F6182, Merck KGaA), whereas NIR samples were mounted in a reducing and oxidizing system (ROXS) buffer containing ascorbic acid as the reducing agent and methylviologen as the oxidizing agent to reduce photobleaching [30]. Samples were sealed with nail polish before imaging.

## 3 Results and discussion

Using the LSEC samples prepared as described above, we acquired super-resolution SIM data sets with OS-SIM, 2D-SIM and enhanced 2D-SIM in both the visible and near-infrared spectral ranges. LSECs are a suitable biological sample for comparing these imaging modalities, as their flat, membrane-dominated structure [31, 32] results in images that provide significant mid-range spatial frequencies, where differences between conventional 2D-SIM and enhanced 2D-SIM become particularly apparent. Furthermore, the small nanopores in the LSEC membranes allow differences in resolution between OS-SIM and the higher-resolution modes to be visualized. Apart from a qualitative image-based comparison of the reconstructed SIM images, we also analysed the different imaging modalities quantitatively in the spatial frequency domain using Power Spectral Density (PSD) and Fourier Ring Correlation (FRC) analyses.

### 3.1 SR-SIM imaging results in the VIS

The results for LSECs stained with BioTracker 555 Orange Cytoplasmic Membrane Dye are shown in Fig. 4. As described in Section 2.1, OS-SIM (Fig. 4(a,d)), 2D-SIM (Fig. 4(b,e)), and enhanced 2D-SIM (Fig.4(c,f)) images were reconstructed from the same raw data set with a fine pattern spacing of 205.7 nm and a coarse pattern spacing of 392.7 nm. All reconstructions were performed with the same reconstruction parameters (NA = 1.5, λ = 595 nm, a = 0.3 for the OTF approximation and 0.9999 in strength (‘a’ value, unitless, normalized) and 20 µm^−1^ FWHM for the OTF attenuation in fairSIM, and the effective pixel size of the reconstructed images is 27.5 nm. In Fig. 4(a-c) we displayed the reconstructed images on the same intensity scale so it becomes immediately visible that the 2D-SIM image shows a reduced signal level. The magnified region of the 2D-SIM image (Fig. 4(e)) is split into two parts with different intensity scales. While the part on the upper left hand side uses the same scaling as the other images, the part on the lower right hand side is rescaled to provide similar contrast to the corresponding regions in the OS-SIM (Fig. 4(d)) and enhanced 2D-SIM (Fig. 4(f)) images. It can be clearly seen that the OS-SIM image has lower spatial resolution compared to the other two images and that the 2D-SIM image is not only showing less overall signal, but also contains honeycomb-style reconstruction artifacts. The artifacts are due to the insufficient suppression of out-of-focus fluorescence and, in extreme cases, these could be misinterpreted as an increased number of cellular nanopores. The resolution enhancement of 2D-SIM and enhanced 2D-SIM compared to OS-SIM is further highlighted by the insets in Fig. 4(d–f), showing nanopores that can only be distinguished in the 2D-SIM and enhanced 2D-SIM data.

**Figure 4:**
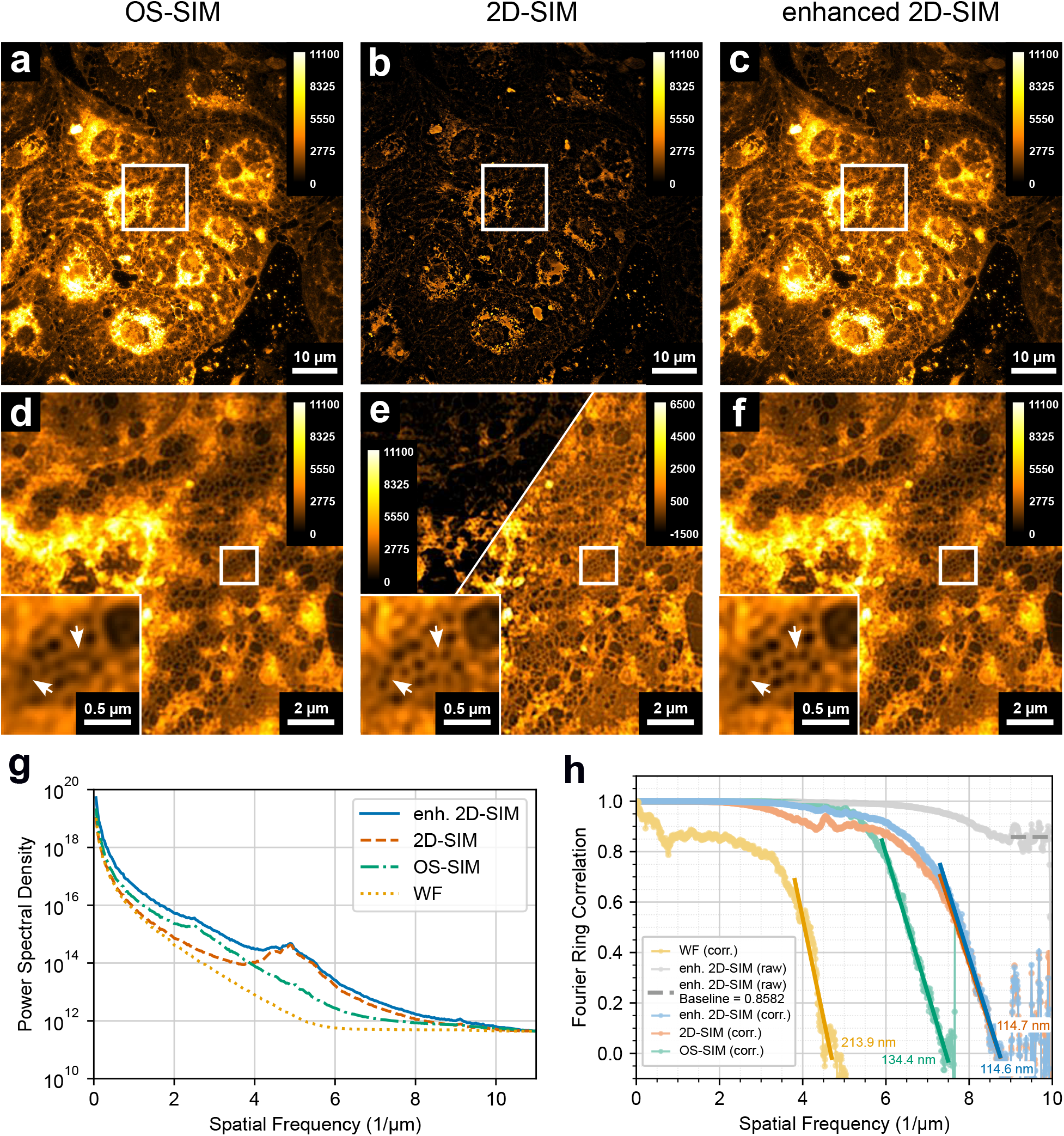
Comparison of imaging performance between OS-SIM, 2D-SIM and enhanced 2D-SIM for membrane-stained LSECs in the VIS. Fixed LSECs membrane-stained with BioTracker 555 were imaged using OS-SIM (a,d), 2D-SIM (b,e) and enhanced 2D-SIM (c,f). Overview images are displayed using the same intensity scale. The 2D-SIM image (b) exhibits a visibly reduced overall signal compared to the OS-SIM (a) and the enhanced 2D-SIM image (c). Magnified regions (d–f) reveal differences in spatial resolution and reconstruction quality. The OS-SIM image shows reduced spatial resolution compared to 2D-SIM and enhanced 2D-SIM, which can be seen in detail for the nanopores in the insets (the intensity scaling was changed to improve the visibility of the nanopores). The 2D-SIM image (e) is split into two parts with different intensity scaling: the part on the upper left hand side uses the same scale as the other images, while the remaining region is rescaled to allow visual comparison to the other images. It can be seen that the 2D-SIM image exhibits more reconstruction artifacts, whereas the enhanced 2D-SIM image (f) shows a smoother membrane and reduced artifact levels, indicating improved reconstruction robustness. (g) Radially averaged power spectral density (PSD) analysis of the reconstructed images reveals complementary frequency support of the different imaging modes. OS-SIM exhibits higher spectral power at low spatial frequencies, reflecting effective background suppression and optical sectioning, while 2D-SIM shows enhanced support at higher spatial frequencies. Enhanced 2D-SIM benefits from both spectral contributions. For visualization purposes, PSD curves were offset by a constant value to align their high-frequency noise plateaus. (h) The corrected FRC curves show that 2D-SIM and enhanced 2D-SIM achieve the same spatial resolution, which is higher compared to OS-SIM. At intermediate spatial frequencies, the FRC curve of 2D-SIM lies below that of OS-SIM, indicating reduced frequency support. Exposure time: 100 ms per single raw SR-SIM image.

The differences between the imaging modalities can further be quantified by the PSD analysis shown in Fig. 4(g). For this analysis, we disabled Wiener-filtering and thus OTF attenuation (see Fig. 2) in fairSIM to obtain fully unfiltered reconstructions. This is necessary as it avoids spectral reshaping by the filters and ensures that the PSD reflects the intrinsic signal and noise transfer characteristics of each imaging mode. Images obtained in this mode are only relevant for quantitative, automated analysis, as without filtering, they, of course, show heavy SIM reconstruction artifacts. Apart from the unfiltered reconstructions of the different imaging modalities, we used the pseudo-widefield image (WF) obtained from fairSIM by averaging all raw SIM image frames as reference. Radially averaged PSDs were calculated from the squared magnitude of the two-dimensional Fourier transforms after subtraction of the mean image intensity to remove the dominant DC component. The Fourier spectra were centered and radially averaged to obtain PSD curves as a function of spatial frequency. At high spatial frequencies, the optical transfer function vanishes and the PSD curves feature a noise-dominated plateau. To facilitate the comparison of spectral support between different imaging modes, the average value of this noise plateau was calculated and used to vertically align the PSD curves such that their high-frequency noise levels coincide with that of the enhanced 2D-SIM reconstruction. This allows for the direct comparison of the spectral shape and relative frequency support between the curves.

The PSD analysis in Fig. 4(g) highlights several distinguishing characteristics between the different imaging modalities. First, the enhanced 2D-SIM PSD curve consistently lies above both, OS-SIM and 2D-SIM curves, indicating improved frequency support across a broad range of spatial frequencies. It is clear that low- and mid-frequency support comes from OS-SIM, while 2D-SIM primarily enhances higher spatial frequencies. The broad maxima in the PSDs around approximately 2.5 µm^−1^ and 5 µm^−1^ arise from the center of the SIM bands (shifted to the corresponding spatial frequencies), where the peaks of each OTF contribute most to the overall signal (see peaks in Fig. 2(b)). Furthermore, the PSD curves show different cut-off frequencies at which they approach the noise plateau, corresponding to the effective spatial resolution of the respective imaging modality. While enhanced 2D-SIM and 2D-SIM have a comparable high cut-off frequency, the PSD curves of OS-SIM and WF reach the noise-dominated regime at substantially lower spatial frequencies. To further quantify the resolution differences, we performed an FRC analysis on the image data shown in Fig. 4(a-c), as presented in Fig. 4(h). FRC is usually performed on two noise-independent acquisitions of the same image [33]. Since only a single raw data set was acquired for each imaging modality, single-image FRC data were calculated [34]. To this end, we applied a coin-flip algorithm to the raw data sets, randomly assigning photon counts to two complementary data sets prior to reconstruction. These two data sets were reconstructed independently and the FRC was calculated from the resulting image pair. To ensure fair comparison between the different imaging modalities, particular care was taken to match the total number of raw frames contributing to each FRC curve. Enhanced 2D-SIM inherently combines raw data from two pattern spacings and therefore uses a larger number of input frames than OS-SIM or conventional 2D-SIM. To avoid differences in noise statistics arising solely from this unequal number of contributing photons, the raw stacks for OS-SIM and 2D-SIM were duplicated prior to application of the coin-flip algorithm, resulting in the same total number of raw frames and thus roughly the same number of photons for each of the imaging modalities. This ensures that differences in the FRC curves reflect reconstruction performance rather than variations in photon statistics or noise averaging. It should be noted, however, that the 2D-SIM data acquired with the finer illumination pattern were recorded prior to the OS-SIM data using the coarser pattern. As a result, mild photobleaching during acquisition cannot be excluded and may have slightly reduced the signal level in the OS-SIM data. Furthermore, we calculated the FRC curve for the WF from Pseudo-WF data generated by averaging all phase images corresponding to one SIM angle. The resulting Pseudo-WF images (one per angle) were used in pairs to calculate FRC curves. The data shown in Fig. 4(h) represent the average of these curves. This was done for the SIM images with the finer pattern, as special care has to be taken when calculating the FRC curve for the WF from the datasets: If the phase images do not average perfectly to the Pseudo-WF image, residual pattern leads to unwanted correlations in the FRC curve. A problem which we observed is a slight phase drift, which is insignificant for the reconstruction process, but causes high auto-correlation in the pseudo-WF data. Using the fine pattern data mitigates this effect somewhat, by shifting the correlation to a higher frequency.

Fixed-pattern noise from the CMOS camera sensor can lead to artificially elevated FRC correlations that exceed what can be physically transmitted by the optical system [13, 35]. Such correlations are not meaningful and should not be part of the FRC analysis for determination of the spatial resolution. We saw these spurious correlations as high-frequency noise plateau in our FRC curves. The FRC data were therefore corrected by fitting the noise plateaus, subtracting their value as a fixed offset from the respective curves, and normalizing the curves to a common noise floor, following a procedure described previously [13]. In accordance with the reference, we have determined the spatial resolution by fitting the linear decay of the corrected FRC curves and calculating the intersection with the fitted noise floor. For clarity, only one representative uncorrected FRC curve of the enhanced 2D-SIM data is shown in Fig. 4(h) in grey.

The FRC analysis displayed in Fig. 4(h) yields a spatial resolution of 114.6 nm for enhanced 2D-SIM, 114.7 nm for conventional 2D-SIM, 134.6 nm for OS-SIM and 213.9 nm for the WF. This is in agreement with the results in Fig. 4(g), confirming that enhanced 2D-SIM and 2D-SIM achieve a comparable spatial resolution that exceeds that of OS-SIM. It should be noted that even the OS-SIM reconstructed images display improved spatial resolution due to a pattern spacing that is comparable with the excitation wavelength after applying the SR-SIM image reconstruction process [27]. The FRC curve of 2D-SIM exhibits an earlier decay at lower spatial frequencies compared to OS-SIM, reflecting the reduced frequency support in the mid-frequency range observed in the PSD analysis. Enhanced 2D-SIM combines the high-frequency resolution of 2D-SIM with improved mid-frequency support.

### 3.2 SR-SIM imaging results in the NIR

To verify the general applicability of this new imaging modality across different systems and wavelength ranges, we repeated the analysis with near-infrared fluorescent samples. The NIR spectral region is of particular interest because it is commonly used for deep-tissue imaging and has different optical transfer characteristics compared to the VIS. Fig. 5 shows the corresponding results for NIR imaging with a fine pattern spacing of 366.2 nm and a coarse pattern spacing of 894.1 nm. OS-SIM (Fig. 5(a,d)), 2D-SIM (Fig. 5(b,e)), and enhanced 2D-SIM (Fig. 5(c,f)) images were reconstructed from the same NIR data set with NA = 1.45, λ = 820 nm, a = 0.1 for the OTF approximation, 0.9999 in strength (‘a’ value, unitless, normalized), and 1.2 µm^−1^ FWHM for the OTF attenuation.The effective pixel size in the NIR experiments was 20.8 nm. As for the VIS data, the 2D-SIM reconstruction exhibits a lower overall signal level compared to OS-SIM and enhanced 2D-SIM, while OS-SIM shows a reduced spatial resolution relative to the other two imaging modes. This can be seen again from the insets in Fig. 5(d–f) where additional nanopores are visible in the 2D-SIM and enhanced 2D-SIM data. A comparison of the magnified regions in Fig. 5(e) and Fig. 5(f) further reveals that the 2D-SIM image contains more reconstruction artifacts than the enhanced 2D-SIM image. This is particularly apparent in the membrane of the LSECs, which appears smoother in the enhanced 2D-SIM reconstruction while maintaining comparable resolution of the nanopores. White arrows in the figures point to regions, where the enhanced spatial resolution can be particularly well seen. The radially averaged PSD analysis for the NIR data (see Fig. 5(g)) shows trends similar to those observed in the visible spectral range (see Fig. 4(g)). The enhanced 2D-SIM curve envelopes the curves from the other imaging modalities, with high-frequency support primarily provided by 2D-SIM and stronger low-to mid-frequency contributions originating from OS-SIM. Compared to the visible measurements, the broad maxima in the PSDs associated with the SIM illumination orders are less pronounced in the NIR, particularly the one at the lower spatial frequency. This can be attributed to the reduced OTF cutoff at longer wavelengths and the generally coarser pattern spacings used for the NIR imaging here, which lead to a stronger overlap of shifted illumination orders with the central frequency region and therefore a smoother curve. Overall, the achievable spatial resolution is lower in the NIR compared to the VIS. This is not only a consequence of the longer excitation and emission wavelengths but also reflects the use of coarser SIM pattern spacings in the NIR imaging experiments. The FRC analysis in Fig. 5(h) yields resolutions of 323.4 nm for the WF, 180.5 nm for enhanced 2D-SIM and 180.0 nm for conventional 2D-SIM, while OS-SIM shows a much lower resolution of 246.5 nm. As in the VIS data, the FRC curve of 2D-SIM exhibits a drop at intermediate spatial frequencies, indicating reduced frequency support in this range.

**Figure 5:**
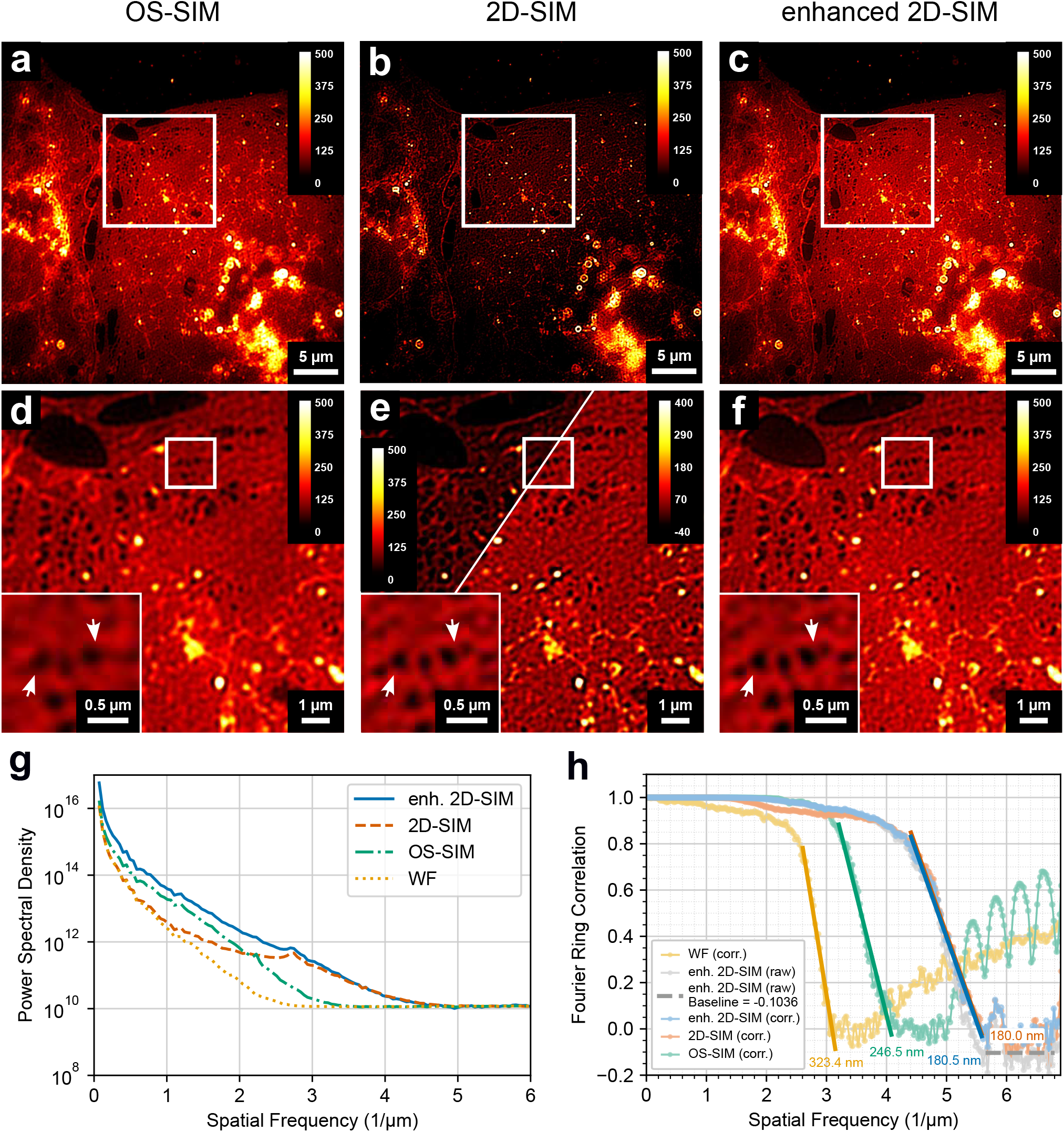
Comparison of imaging performance between OS-SIM, 2D-SIM and enhanced 2D-SIM for membrane-stained LSECs in the NIR. Fixed membrane-stained LSECs labeled with BioTracker NIR790 were imaged using OS-SIM (a,d), 2D-SIM (b,e) and enhanced 2D-SIM (c,f). The overview images (a–c) show a FOV of 38.1 µm × 38.1 µm and are displayed using the same intensity scale. The 2D-SIM image (b) appears much darker than the other images due to its reduced overall signal level. Magnified regions reveal that OS-SIM (d) exhibits lower spatial resolution, while enhanced 2D-SIM (f) shows fewer reconstruction artifacts compared to 2D-SIM (e). This is particularly visible on the membrane of the LSECs, which appears smoother in the enhanced 2D-SIM image as a result of improved support in the mid-frequency range. The arrows in the insets point to regions where the resolution enhancement of 2D-SIM and enhanced 2D-SIM compared to OS-SIM is apparent. The insets were scaled in intensity to visualized the differences between the images better. The effective pixel size is 20.8 nm. The radially averaged PSD analysis (g) supports these observations. While high-frequency support in enhanced 2D-SIM originates primarily from the 2D-SIM data, OS-SIM dominates the low-to mid-frequency range. The different cut-off frequencies reflect the differing spatial resolutions of the imaging modes. The corrected FRC curves (h) show that enhanced 2D-SIM and conventional 2D-SIM achieve the same spatial resolution, whereas OS-SIM reaches a lower spatial resolution. In addition, the 2D-SIM FRC curve exhibits a drop at intermediate spatial frequencies. For clarity, only one representative uncorrected FRC curve from the enhanced 2D-SIM data is shown. Exposure time: 100 ms per single raw SR-SIM image.

In summary, the NIR results reproduce the same characteristic differences between the imaging modalities observed in the VIS and demonstrate that enhanced 2D-SIM provides a robust combination of background suppression and frequency support across different wavelength ranges.

## 4 Conclusion

We have presented a novel 2D-SIM imaging approach termed *enhanced 2D-SIM*, which combines the complementary strengths of OS-SIM and conventional 2D-SIM within a single image reconstruction process. By acquiring raw data using both, fine and coarse illumination patterns, enhanced 2D-SIM enables the reconstruction of super-resolved fluorescence images with reduced reconstruction artifacts and increased suppression of out-of-focus light. Besides the theoretical framework underlying this method based on the comparison of optical transfer functions, we have demonstrated the practical applicability of enhanced 2D-SIM in the VIS and NIR spectral ranges for imaging transcellular nanopores in the plasma membrane of LSECs. The differences between enhanced 2D-SIM, 2D-SIM and OS-SIM were characterized through direct comparison of the image data as well as quantitative analyses based on PSD and FRC. It was shown that enhanced 2D-SIM achieves the same spatial resolution as 2D-SIM while providing an improved overall signal level and reduced reconstruction artifacts. This is based on the improved frequency support of enhanced 2D-SIM where high-frequency support is gained from 2D-SIM and low-to mid-frequency support from OS-SIM raw image data.

The enhanced 2D-SIM approach is wavelength-independent and can easily be applied to existing 2D-SIM (and 3D-SIM) setups as long as they have the functionality to switch between a coarser and finer SIM pattern. Since in practice, 3D-SIM is a far more challenging method to implement in optical systems compared to 2D-SIM modalities, this enhanced 2D-SIM approach is a practical alternative for 2D-SIM systems where good lateral spatial resolution, reconstruction stability and background suppression is needed.

## Acknowledgements

Funding for this project was provided by the European Union’s European Innovation Council PATH-FINDER Open Programme under grant agreement No 101046928. H.O. and M.M. received funding from the European Regional Development Fund (EFRE/JTF-Programm NRW 2021-2027), funding line “Start-up Transfer.NRW” under the project “ProSIM” and the EXIST-Research Transfer program (Phase I) of the German Federal Ministry for Economic Affairs and Energy under grant No 03EFNW0436. S.M.S. was, in the early stages of this project, funded by the German Federal Ministry for Research, Technology and Aeronautics (BMFTR), through project BetterView (FKZ 13N15827, 13N15830). T.H. and J.S.a.E. further acknowledge funding by the Deutsche Forschungs-gemeinschaft (DFG, German Research Foundation), through grant no. 540217954. Open Access funding is enabled and organized by Projekt DEAL. The authors would like to thank Jochen Linnenbrügger for the CAD design of the HWP rotator and the VIS-SIM setup. We would also like to thank Sara Abrahamsson, Reto Fiolka and Rainer Heintzmann for fruitful discussions that inspired the project.

## Data Availability Statement

Raw data for both microscopy figures (Fig. 4,5) is publicly available on zenodo (DOI 10.5281/zenodo.18760753).

## Author contributions

S.M.S. built the NIR imaging system, acquired the NIR imaging data, performed data analysis, and drafted the manuscript. H.O. built the VIS imaging system and acquired the VIS data. J.C.S.-S., A.K. and A. K.-K. prepared the biological samples. M.M. conceived the idea for this study and performed the OTF calculations. J.S.a.E. and T.H. provided funding and supervised the work. All authors reviewed and approved the final manuscript.

